# Site-specific quantification of the *in vivo* UFMylome reveals myosin modification in ALS

**DOI:** 10.1101/2024.10.30.621144

**Authors:** Ronnie Blazev, Barry M. Zee, Hayley Peckham, Yaan-Kit Ng, Christopher T. A. Lewis, Chengxin Zhang, James W. McNamara, Craig A. Goodman, Paul Gregorevic, Julien Ochala, Frederik J. Steyn, Shyuan T. Ngo, Matthew P. Stokes, Benjamin L. Parker

**Affiliations:** Department of Anatomy and Physiology, The University of Melbourne, VIC, Australia; Centre for Muscle Research, The University of Melbourne, VIC, Australia; Cell Signaling Technology, Danvers, MA, USA; Department of Biomedical Sciences, University of Copenhagen, Copenhagen, Denmark; Novo Nordisk A/S, Research and Early Development, Måløv, Denmark; Department of Computational Medicine and Bioinformatics, University of Michigan, Ann Arbor, MI, USA; Murdoch Children’s Research Institute and Melbourne Centre for Cardiovascular Genomics and Regenerative Medicine, The Royal Children’s Hospital, Parkville, VIC, Australia; Melbourne Centre for Cardiovascular Genomics and Regenerative Medicine, The Royal Children’s Hospital, Melbourne, VIC, Australia; Novo Nordisk Foundation Center for Stem Cell Medicine, Murdoch Children’s Research Institute, Melbourne, VIC, Australia; Department of Neurology, University of Washington School of Medicine, Seattle, WA, USA; Department of Neurology, Royal Brisbane and Women’s Hospital, Brisbane, QLD, Australia; School of Biomedical Sciences, The University of Queensland, Brisbane, QLD, Australia; Australian Institute for Bioengineering and Nanotechnology, The University of Queensland, Brisbane, QLD, Australia; Centre for Clinical Research, The University of Queensland, Brisbane, QLD, Australia

**Keywords:** Ubiquitin Fold Modifier 1 (UFM1), UFMylation, ubiquitin-like modification

## Abstract

UFMylation is a ubiquitin-like protein post-translational modification of Ubiquitin Fold Modifier 1 (UFM1) applied to substrate proteins. The UFMylation system is important for normal development and plays a critical role in a variety of cellular processes including regulating telomere length, stress responses and protein quality control. Here, we describe the development of an antibody-based enrichment approach to immunoprecipitate *in vivo* remnant UFMylated peptides and identification by liquid chromatography and tandem mass spectrometry (LC-MS/MS). We used this approach to identify >200 UFMylation sites from various mouse tissues revealing extensive modification in skeletal muscle. Furthermore, we show that UFMylation is increased in skeletal muscle biopsies from people living with Amyotrophic Lateral Sclerosis (plwALS). Quantification of UFMylation sites in these participant biopsies with multiplexed isotopic labeling and LC-MS/MS reveal prominent increases in myosin UFMylation. Finally, *in silico* modelling suggest UFMylation of myosin directly adjacent to the ATP-binding site may regulate stability and/or function. Our data suggest that although UFMylation is not as widespread as ubiquitylation, its *in vivo* status is more complex than previously thought.

## Introduction

The Ubiquitin Fold Modifier 1 (UFM1) covalently modifies Lys residues of substrate proteins in a process called UFMylation and has been implicated in various cellular processes and diseases (1, 2). UFMylation begins with removal of the C-terminal Ser84Cys85 residues of UFM1 by UFSP1/2 to generate UFM1ΔSC which is subsequently activated by forming a thioester bond with UBA5, a UFM1-specific E1 activating enzyme. Next, activated-UFM1ΔSC is transferred to the UFM1-specific conjugating E2 enzyme, UFC1. Together with the UFM1-specific ligase E3 complex (UFL1, DDRGK1 and CDK5RAP3), UFM1ΔSC is then transferred to substrate Lys residues (3, 4). The Lys residues of conjugated-UFM1 can then undergo further poly-UFMylation (5, 6). Currently, <15 UFMylation substrates have been identified primarily via expression of exogenously epitope-tagged UFM1 and enrichment from cell culture models (for a summary see (2)). Here, we describe a new enrichment method coupled to LC-MS/MS to site-specifically identify and quantify endogenous UFMylation sites *in vivo*. We applied this method to quantify UFMylation sites in human skeletal muscle biopsies from plwALS participants and heathy age-matched controls revealing extensive changes in myosin UFMylation.

## Results

We generated an antibody-based enrichment strategy coupled to LC-MS/MS to identify *in vivo* UFMylation sites. The approach is based on the identification of ubiquitination sites using tryptic peptide immunoprecipitation with the “remnant diGly” anti-GG-ε-K antibody (7). Briefly, following generation of UFM1ΔSC, the sequential action of UFMylation E1-E2-E3 ligases transfer UFM1ΔSC to substrate Lys residues within cells (**Fig. 1A**). Proteins are extracted with denaturing buffers and proteolytically digested with trypsin which cleaves C-terminal to Lys and Arg residues. Trypsin cleaves after Arg81 on UFM1ΔSC- conjugated substrate proteins and leaves a “remnant” ValGly attached to the substrate Lys via an isopeptide bond (VG-ε-K isopeptide). We generated three monoclonal pan anti-VG-ε-K antibody clones to immunoprecipitate “remnant VG” UFMylated sites independently from the surrounding amino acid context. ELISA showed the antibody clones had ∼6-17-fold enhanced specificity to VG-ε-K- compared to GG-ε-K-containing peptides (**Fig. 1B**). To screen these antibody clones, tryptic peptides from mouse *gastrocnemius* skeletal muscles were immunoprecipitated with each of the three variants or a pooled cocktail followed by 2D-LC-MS/MS and analysis with two different search algorithms (Sequest+Percolator and MSFragger+PTM-Prophet). In total, we identified 385 unique VG-modified peptides with either search algorithm with each clone identifying unique subsets, and the pooled cocktail identifying the greatest number (**Fig. 1C-E**). Motif analysis identified no major sequence specificity but a slight upstream acidic amino acid residue preference for all clones where it was most prominent for clone 2 (**Fig. 1F**). A total of 199 unique VG-modified peptides were identified by both algorithms representing the greatest confidence (**Fig. 1G**)(**Table S1**). This includes the previously identified and independently validated UFMylation site of Lys134 on RPL26 (6, 8). Because a high proportion of the skeletal muscle proteome comprises of contractile proteins, we sought to test the capacity of our methods to identify UFMylation of other substrates across other tissue and cell types. Comparing global UFMylation abundance using anti-UFM1 western blot in various mouse and human cells, and in various mouse tissues revealed consistent and prominent bands at ∼30 and 60 kDa (**Fig. 1H**). However, the tissue samples exhibited increased immunoreactivity across a range of molecular weights suggesting more complex UFMylation *in vivo* including tissue-specific substrates. We next performed tryptic peptide immunoprecipitations in these mouse tissues using the pooled anti-VG-ε-K antibody cocktail followed by 2D-LC-MS/MS and considered only data identified by both search algorithms (**Table S2**). **Fig. 1I** plots the number of VG-containing peptide spectral matches (PSMs), unique peptides and sites identified by both search algorithms for each tissue. A total of 250 unique VG-containing peptides were identified and enrichment analysis identified over-representation of proteins associated with the contractile apparatus, or localized to extracellular, endoplasmic/sarcoplasmic reticulum (ER/SR) or the mitochondrial Gene Ontology (GO) compartments (**Fig. 1J**). RPL26 UFMylation was identified across multiple tissues but other previously identified sites on histone H4 (9), p53 (10), ASC1 (5), CYB5R3 (11) and DDRGK1 (3) were not identified. KEGG pathway analysis revealed UFMylated proteins were associated with central carbon metabolism, amino acid/glucose metabolism and proteins involved in muscle contraction/cardiomyopathy (**Fig. 1K**). Analysis of UFMylated proteins forming interaction networks in the STRINGdb identified inter-connected associations of the contractile apparatus, proteins involved in calcium handling at the ER/SR, glucose metabolism and translational regulators (**Fig. 1L**). We also identified VG modification of Lys19 and Lys69 on UFM1 providing direct evidence for the presence of di- or poly-UFMylation (5, 6).

**Figure 1.**
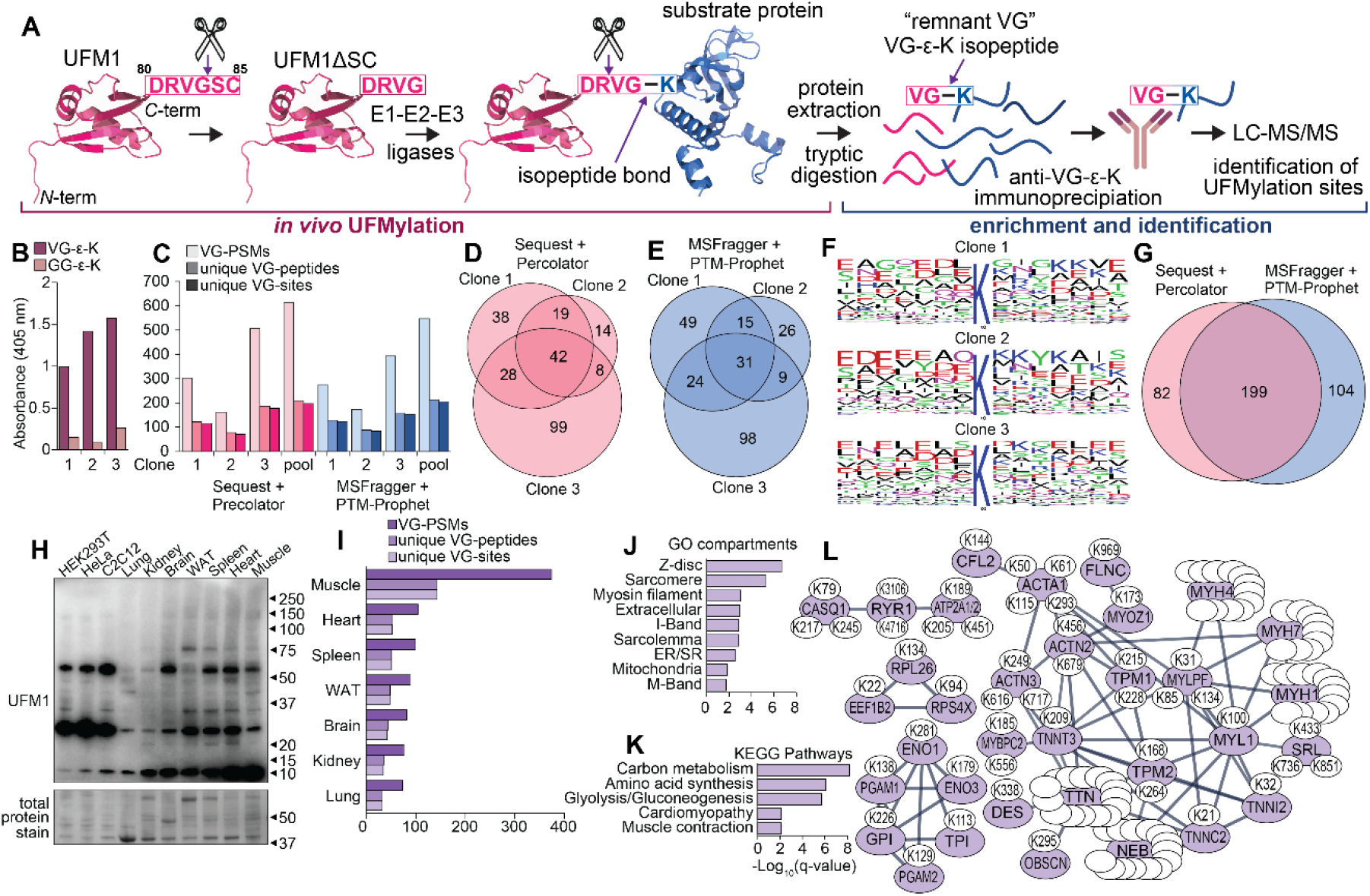
Enrichment and site-specific identification of the UFMylome. (**A**) Overview of UFMylation and schematic for the enrichment of VG-modified peptides and identification by liquid chromatography tandem mass spectrometry (LC-MS/MS). (**B**) Analysis of antibody specificity to VG-ε-K- compared to GG- ε-K-containing peptides by ELISA. (**C**) Number of VG-peptide spectral matches (PSMs) and unique VG- peptides/VG-sites identified by each of the antibody clones or the pooled cocktail. (**D-E**) Overlap of VG- peptides identified by each antibody clone. (**F**) Motif enrichment of the amino acids surrounding the UFMylation sites. (**G**) Overlap of VG-peptides identified by each search algorithm. (**H**) Anti-UFM1 western blot of various mouse and human cell lines, and various mouse tissues. (**I**) Number of VG-peptide spectral matches (PSMs) and unique VG-peptides/VG-sites identified in each mouse tissue. (**J**) Gene Ontology (GO), and (**K**) Kyoto Encyclopedia of Genes and Genomes (KEGG) enrichment analysis of the UFMylated proteins. (**L**) STRINGdb analysis of the UFMylated proteins.

We next sought to validate a subset of skeletal muscle UFMylation sites by knocking down UFC1 which should result in concomitant decrease in their abundance. Mouse *extensor digitorum longus* muscles were injected with recombinant adeno-associated virus serotype 6 (rAAV6) with the left leg receiving scramble negative control shRNA and the right contralateral leg receiving UFC1 shRNA’s. Muscles were harvested after 28 days and western blotting revealed a decrease in UFC1 protein levels with a concomitant decrease in global UFMylation (**Fig. 2A**). Tryptic peptides were immunoprecipitated using pooled anti-VG-ε-K antibody clones followed by multiplex stable isotope labelling with Tandem Mass Tags (TMT) and analysis by 2D-LC-MS/MS. A total of 22 unique VG-containing peptides were quantified in every sample and a general decrease in UFMylation was observed following knockdown of UFC1 (**Fig. 2B**)(**Table S3**). The intensity of VG-containing peptides was normalized to total protein levels obtained via a parallel proteomic analysis without anti-VG-ε-K enrichment. This identified 10 down-regulated VG- containing peptides at a q<0.05 using Benjamini-Hochberg false discovery rate while 12 were down- regulated at a less strict p<0.05. The MS/MS spectra of the most down-regulated peptides were manually annotated and mapped to multiple myosin isoforms given their shared sequence similarity such as Lys1381/1381/1375/1378/1381 on MYH1/2/3/4, respectively; Lys975/975/969/972 on MYH1/2/3/4, respectively; and 1795/4 on MYH4/8, respectively (**Fig. 2C-E**). The fewer VG-containing peptides identified in these knockdown experiments compared to the data described above is likely due to the >10- fold less available starting material (100 µg / sample) combined with the reduced identification efficiency of highly charged TMT-labelled peptides (12). Despite this, our data validate that myosin is UFMylated *in vivo*.

**Figure 2.**
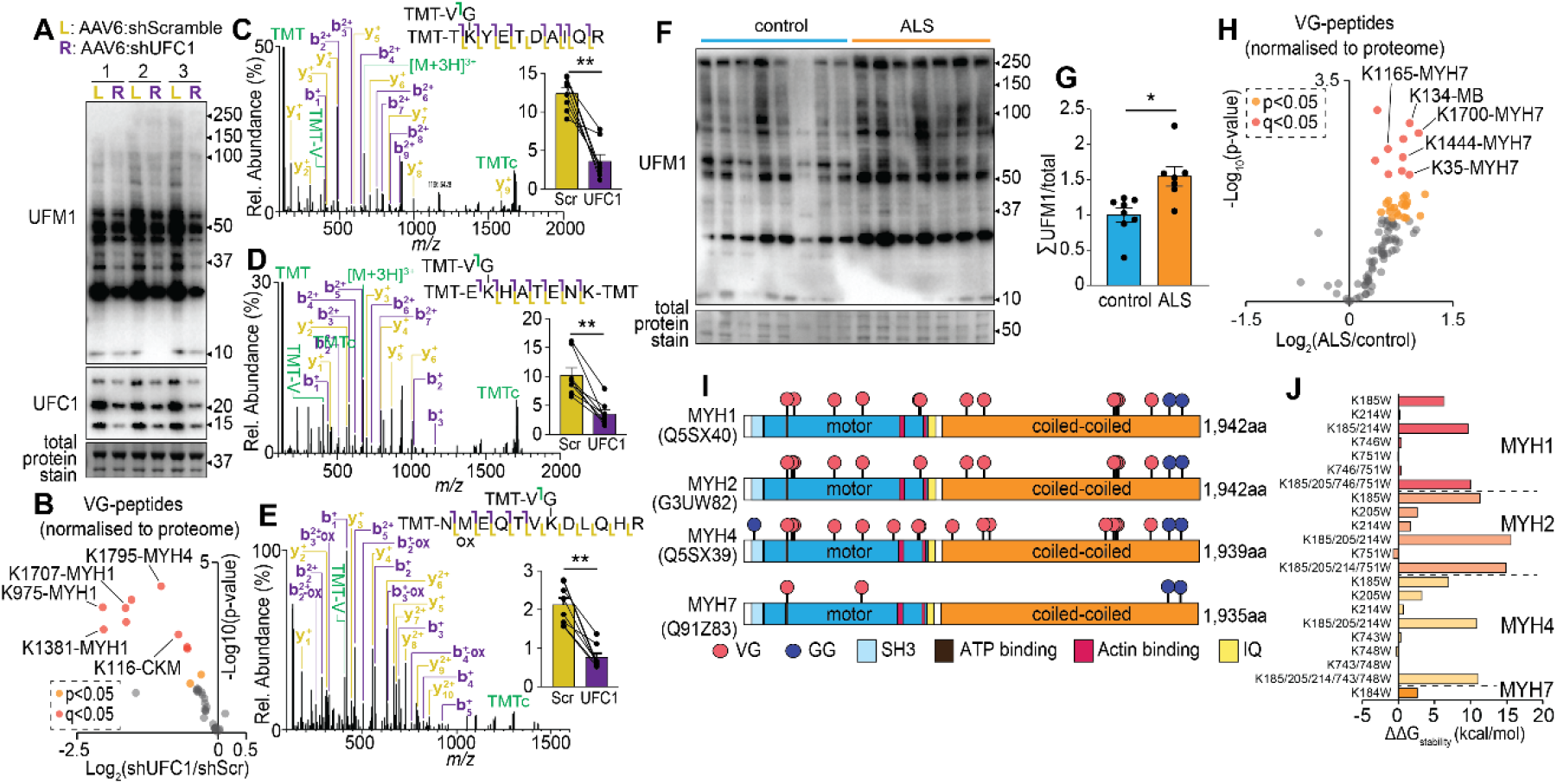
Site-specific quantification of the UFMylome identifies a differential signature in human skeletal muscle associated with ALS, and specific sites of UFMylation on myosin proteins. (**A**) Anti-UFM1 western blot of mouse *extensor digitorum longus* (EDL) skeletal muscles treated with recombinant adeno-associated virus serotype 6 (rAAV6) harboring scramble negative control shRNA (L: left leg) or UFC1 shRNA (R: right leg). (**B**) Volcano plot of quantified VG-peptides from mouse EDL following knockdown of UFC1. Annotated MS/MS of VG-containing peptides from (**C**) Lys1381/1381/1375/1378/1381 on MYH1/2/3/4, (**D**) Lys975/975/969/972 on MYH1/2/3/4, and (**E**) 1795/4 on MYH4/8, respectively. Inserts show Log2(Area under the curve) normalized to total myosin levels. ^**^q<0.01; paired student t-test with Benjamini Hochberg FDR. Error bars: SEM. (**F**) Anti-UFM1 western blot of skeletal muscle biopsies from individuals diagnosed with ALS and age-matched controls, and (**G**) Immunoreactivity densitometry of the entire lane. ^*^p<0.05; unpaired student t-test. Error bars: SEM. (**H**) Volcano plot of quantified VG-peptides from human individuals. (**I**) VG- and GG-peptides identified on myosin purified from mouse skeletal muscle. (**J**) Modelling the effects of Lys>Trp substitutions on myosin stability.

Skeletal muscle UFMylation has been shown to increase in the SOD1(G37R) mouse model of ALS (13). To investigate if this is also observed in humans, we analyzed biopsies of *vastus lateralis* skeletal muscle from plwALS and age-matched controls, and previously shown that these plwALS display disease-associated atrophy (14). Western blotting revealed a conserved increase in the UFMylation of several proteins (**Fig. 2F-G**). To identify and quantify these UFMylation sites, tryptic peptides were immunoprecipitated from each of the participants’ biopsies using pooled anti-VG-ε-K antibody clones followed by TMT labelling and analysis by 2D-LC-MS/MS. A total of 56 unique VG-containing peptides were quantified in every biopsy and a general increase in UFMylation was observed in plwALS (**Fig. 2H**)(**Table S4**). Nine unique VG-containing peptides were up-regulated at a q<0.05 using Benjamini-Hochberg false discovery rate while 24 were up-regulated at a less strict p<0.05. The increase in abundance of these VG-containing peptides was observed independent of changes in their corresponding protein abundance suggesting site-specific changes in UFMylation. The most significantly up-regulated UFMylation sites were observed on the contractile protein Conventional Myosin-7 (MYH7). Other up- regulated UFMylation sites at the less stringent cut-off were on other contractile and/or structural proteins such as Troponin C1 (TNNC1) and Titin (TNN). To provide a more detailed analysis of UFMylation and potential interplay with ubiquitinylation, we next purified myosin isoforms from mouse *tibialis anterior* muscle and analyzed tryptic peptides by LC-MS/MS. Twenty unique VG-containing peptides and four unique GG-containing peptides were identified (**Table S5**). The majority of these VG-/GG-containing peptides mapped to multiple myosin isoforms given their shared sequence similarity. Mapping VG- and GG-sites onto the primary amino acid sequences of the major isoforms of MYH1, MYH2, MYH4 and MYH7 revealed several interesting clusters (**Fig. 2I**). For example, Lys185 and 214 on MYH1/2/4, Lys 204 on MYH2/4, and Lys184 on MYH7 which are directly adjacent to the ATP binding site. Other interesting sites include Lys746 on MYH1, Lys743 on MYH4, and Lys751 on MYH1/2 which are between the two actin-binding domains. To investigate potential functional impacts, we used EvoEF, an *in silico* modelling approach to estimate the effects of single amino acid substitutions on protein stability using physical energy function (15). Here, Lys residues were mutated to Trp either in isolation or in combinations. Mutation of residues adjacent to the ATP binding site dramatically increased ΔΔG_Stability_ of >10 kcal/mol while the sites between the two actin-binding domains had little impact (**Fig 2J**). For context, the observed ΔΔG_Stability_ changes constitute a >10-fold increase in predicted stability compared to myosin phosphorylation (16). Although this *in silico* approach does not directly assess the functional impact of myosin UFMylation, it highlights that modification of specific amino acids are highly likely to alter myosin the function. Taken together, our site-specific identification of UFMylation has validated myosin as a target of this modification that is up-regulated during ALS in human patients.

## Discussion

To date, knowledge of how UFMylation influences the proteome has been limited by challenges associated with identifying proteins specifically targeted by the UFMylation system. To address this limitation, developed an approach to identify and quantify the endogenous UFMylome *in vivo* using tryptic peptide immunoprecipitation using pan anti-VG-ε-K antibodies coupled with multiplexed stable isotope labelling and LC-MS/MS. This approach provides distinct advantages over traditional approaches to identify UFMylated substrates which rely on the expression of exogenously epitope-tagged UFM1 followed by protein-level enrichment. Firstly, peptide-level enrichment may increase the ability of the mass spectrometer to sequence the VG-containing peptides to pinpoint the modification sites. Secondly, samples can be lysed in very harsh denaturing conditions to inhibit all enzyme activity, whereas these lysis buffers may not be compatible with epitope-tagged protein-level enrichment. Additionally, our approach avoids the need for the over-expression of exogenously epitope-tagged UFM1 to un-naturally high concentrations that may result in non-physiological substrate UFMylation. This phenomenon has previously been observed with high levels of ubiquitin leading to ubiquitylation of non-physiological substrates (17, 18). It is worth noting that several previous studies identifying UFMylated substrates have also combined ectopic expression of tagged-UFM1 with genetic manipulations of the E1-E2-E3 ligase system to further boost cellular UFMylation levels.

Our data have identified more than 200 novel UFMylation sites and reveal that this modification is more widespread than previously thought. The UFMylation machinery is enriched at the ER, and CRISPR screens have identified a key role of this modification in various ER stress responses (6, 19). A principal mechanism is via the UFMylation of RPL26 and the regulation of ribosome-associated quality control in stalled translocon (8, 20, 21). In addition to RPL26, we identified other UFMylated proteins enriched at the ER/SR including CASQ1, RYR1 and ATP2A1/2 which play critical roles in calcium handling. As UFMylation negatively regulates skeletal muscle contractile function (13), and given we observed UFMylation of both calcium handling proteins and the contractile apparatus particularly myosin, future experiments have the opportunity to delineate the exact role of this modification upon the excitation-contraction pathway. The extensive modification of myosin also warrants further investigation particularly sites adjacent to the ATP binding domain which given our *in silico* predictions suggest large changes in stability following mutation of these amino acids. Furthermore, whether the activation of UFMylation is causal or protective in diseases such as ALS warrants further investigation.

While our approach for identification and quantification of substrate UFMylation offer specific advantages, a potential limitation of our VG-peptide-level enrichment and identification of ‘remnant’ UFMylation is that the required trypsin cleavage results in loss of information on site-specific polyUFMylation or linkage patterns. As such, we envisage that considered application of our techniques will help to expand understanding of UFMylation across cell types and biological contexts, but that ongoing investigation of opportunities to develop complementary methods hold the potential to further expand insight into the full scale of UFMylation complexity.

## Materials and Methods

### Experimental details are provided in SI Appendix

#### Anti-VG-ε-K antibody production and bead production

New Zealand White rabbits were immunized with a KLH-conjugated degenerate peptide library containing a VG-remnant modified lysine. Serum samples were screened for PTM specificity by ELISA.

#### Anti-VG-ε-K peptide enrichment

Tryptic peptides were resuspended in 500 µl of Immunoprecipitation buffer (50mM MOPS, 10mM NaH2PO4, 50mM NaCl, pH 7.5), and adjusted to a pH of 7.5. Peptides were rotated with antibody:Protein-G bead complex overnight at 4°C.

## Data availability

The mass spectrometry proteomics data have been deposited to the ProteomeXchange Consortium via the PRIDE (15) partner repository with the dataset identifier PXD051412 (Username; Password: 3se7c5iG).

## Acknowledgments

We thank Nicholas Williamson, Ching-Seng Ang, Shuai Nie, Swati Varshney and Michael Leeming for instrument support in the Bio21 Mass Spectrometry and Proteomics Facility. We thank all patients with ALS and control individuals who participated in this study. This work was funded by an Emerging Leader Investigator Grant (APP2009642) from the National Health and Medical Research Council (NHMRC)(Australia) and a University of Melbourne Driving Research Momentum Grant to B.L.P., a Motor Neurone Disease Research Australia Charcot Grant and NHMRC Ideas Grant 1185427 to F.J.S. and S.T.N., and the Scott Sullivan MND Research Fellowship (University of Queensland, Royal Brisbane & Women’s Hospital, and the MND and Me Foundation) to S.T.N and a NHMRC Investigator Grant (APP2017070) to P.G. This work was also supported by a Department of Anatomy and Physiology (The University of Melbourne) ECR Seeding Grant to R.B. Y.-K.N. is a recipient of a School of Biomedical Science Postgraduate Award.

