## Supporting Information for "Site-specific quantification of the *in vivo* UFMylome reveals myosin modification in ALS"

**Materials and Methods**

**Cell culture**

C2C12, HEK293T and HeLa cells were grown in Dulbecco's Modified Eagle Medium (DMEM) (GIBCO by Life Technologies; # 11995065), supplemented with 10% fetal bovine serum (FBS) (Life Technologies; #26140079), pyruvate and GlutaMAX (GIBCO by Life Technologies). Cells were kept at 37°C and 5% CO2 in a humidified incubator Direct Heat CO2 Incubator featuring Oxygen Control (In Vitro Technologies).

**Mouse housing and rAAV6 intramuscular injection**

All mouse experiments were approved by The University of Melbourne Animal Ethics Committee (#26527), and adhere to the guidelines set out by the National Health and Medical Research Council of Australia for the care and use of animals for scientific purposes. Male C57BL/6J mice were housed at 22°C (±1°C) in groups of five/cage and maintained on a Standard Chow diet (Specialty Feeds, Glen Forest, WA, Australia, #SF15-049) with a 12-hr light/dark cycle and ad libitum access to food and water. rAAV6 production, purification and shRNA sequences are previously described (1). For intramuscular injections of rAAV6, mice were anesthetized with isoflurane (4% in oxygen at 1L/min) and then transferred to a dissecting microscope stage with heat pad and isoflurane inhalation nose piece (2% in oxygen at 1 L/min). Unconsciousness was assessed via the lack of leg and optical reflexes for at least one minute to ensure head position does not affect normal breathing. Mice received subcutaneous analgesic injection of meloxicam between the shoulder blades (5mg/kg) and the surface of the hindlimbs were sterilized with 80% ethanol. The EDL muscles were injected with 2 x 10^10^ vector genomes / 30 µl of rAAV6 using a 32G needle. Mice were returned to cages and body weights monitored daily for the first 3 days and then weekly prior to cull at 28 days post-injection.

**Human ALS participants and experimental procedures**

The human participants, biopsy collection and lysates were generated as described previously (2). Patient characteristics and ethics approval is provided in this previous publication and the current study analysed the identical participant biopsy lysates.

**Anti-VG-ε-K antibody production and bead production**

New Zealand White rabbits were immunized with a KLH-conjugated degenerate peptide library containing a VG-remnant modified lysine; serum samples were screened for PTM specificity by ELISA. Monoclonal antibodies were then generated using proprietary methods developed at Cell Signaling Technology, Danvers, MA. Each IP was performed with 1:10 antibody:protein ratio e.g 1mg of antibody:10mg of digested mouse protein lysate or 100 µg of antibody:1 mg of digested human muscle biopsy protein lysate. Antibodies were incubated on Protein-A agarose beads at a ratio of 100 µg of antibody / 30 µl of beads overnight at 4°C with rotation. The beads were washed four times with 1 ml of PBS and used immediately for the enrichment. The pooled cocktail mix was generated by conjugating each antibody clone and then pooling the beads. For comparison of antibody clones vs the pooled antibody cocktail, the amount of antibody on the beads was constant e.g 1 mg for individual clones vs 3 x 333 µg of individual clones mixed together.

**Lysate preparation**

Tissue was lysed by tip-probe sonication in 6M guanidine HCl (Sigma-Aldrich, St. Louis, MO, USA, #G4505), 100mM Tris pH 8.5 containing 10mM Tris(2-carboxyethyl)phosphine (Sigma-Aldrich, #75259) and 40mM 2-Chloroacetamide (Sigma-Aldrich, #22790). The lysate was then heated at 95°C for 5 mins and centrifuged at 18,000 g for 30 mins at 4°C. The supernatant was diluted 1:1 with LC-MS/MS water and precipitated overnight in a final concentration of 80% acetone at -30°C. The lysate was centrifuged at 4,500 g for 5 mins at 4°C and the supernatant was disregarded. The protein pellet was washed with 80% ice-cold acetone and centrifuged at 4,500 g for 5 mins at 4°C, then resuspended in digestion buffer (10% trifluoroethanol (Thermo Fisher Scientific, #96924) in 100 mM HEPES pH 7.5). Protein was quantified using a BCA protein assay (Thermo Fisher Scientific, #23325) and normalized to 2µg/µl containing Laemelli buffer (BioRad, Hercules, CA, USA, #1610747) for western blotting. For comparison of the 3 antibody clones and pooled cocktail, 4 x 10 mg protein aliquots of the identical mouse skeletal muscle was normalized to 1 ml each. For enrichment of VG-peptides from mouse tissues, 10mg of protein was normalized to 1 ml of digestion buffer, while for enrichment from human skeletal muscle biopsies, 1mg of protein was normalized to 200 µl. Protein lysates were digested in sequencing grade trypsin (Sigma-Aldrich, #T6567) and LysC (Wako, Chuo-Ku, Osaka, Japan, #129-02541) (1:100 enzyme:substrate ratio) overnight at 37°C with shaking at 2000 rpm.

**Western blotting**

Proteins were separated on NuPAGE 4-12% Bis-Tris protein gels (Thermoscientific) in MOPS SDS running buffer at 145V for 60 mins at room temperature. Proteins were then transferred to PVDF membranes (Millipore, Burlington, MA, USA, #IPFL00010) in NuPAGE transfer buffer at 20V for 60 mins at room temperature. Membranes were blocked with 5% skim milk in Tris-buffered saline containing 0.1% Tween-20 (TBST) for at least 60 mins at room temperature with gentle shaking. Membranes were then incubated overnight at 4°C with gentle shaking in anti-UFM1 antibody (1:1000) (Abcam; ab109305) containing 5% BSA, 0.02% NaN3 in TBST. Membranes were incubated with HRP-secondary antibody in 5% skim milk in TBST for at least 60 mins at room temperature with gentle shaking. Immunoreactivity was visualized with Immobilon Western Chemiluminescent HRP Substrate (Millipore, #WBKLS0500) and imaged on a ChemiDoc (BioRad). Membranes were stained with BLOt-FastStain (GBiosciences; 786-34) for protein visualization. Densitometry was performed in Image J (3).

**Peptide purification and anti-VG-ε-K enrichment**

Peptides were acidified by Trifluoroacetic acid (TFA) (Thermo Fisher Scientific, #FSBA116) to a final concentration of 1% TFA and centrifuged at 16,000 g for 10 mins at room temperature. Peptides were desalted using reversed-phase chromatography with HLB-SPE columns (Waters, Milford, MA, USA) and washed with 0.1% TFA, then eluted with 80% acetonitrile, containing 0.1% TFA. Peptides were vacuum-dried at 45°C. Peptides were resuspended in 500 µl of Immunoprecipitation buffer (50mM MOPS, 10mM NaH2PO4, 50mM NaCl, pH 7.5), and adjusted to a pH of 7.5 with 5M NaOH, then centrifuged at 16,000 g for 5 mins at 4°C. The supernatant containing peptides were rotated with antibody:Protein-G bead complex overnight at 4°C. The beads were washed three times with 1ml of IP buffer and then twice with 1 ml of LC-MS/MS water. Peptides were eluted twice with 75 µl of 0.2% TFA for 5 mins at 10°C with shaking at 1200 rpm, and beads filtered using in-house packed C8 tips (3M, Saint Paul, MN, USA, #11913614). Enriched peptides from mouse tissues were purified through in-house packed SDB-RPS columns as described below. Enriched peptides from human skeletal muscle lysates were dried by vacuum centrifugation and resuspended in 10 µl 200mM HEPES, pH 7.4, for 5 mins with shaking at 2000 RPM, and then 80 µg/10 µl of TMTpro-16plex in 100% acetontrile was added, followed by 1 hr incubation at room temperature (Thermo Fisher Scientific, #A44520). The labelling scheme and sample identification has been uploaded to the PRIDE Proteomics repository as described below. The reaction was deacylated to a final concentration of 0.3% hydroxylamine then quenched with 1% TFA. All 16 samples were pooled together and purified through in-house packed SDB-RPS (Sigma-Aldrich, #66886-U) tip, and washed with 99% isopropanol, 1% TFA followed by 5% acetonitrile, 0.2% TFA, then eluted with 80% acetonitrile, 5% NH4OH. Sample was resuspended in 2% acetonitrile, 0.1% TFA and stored at -80°C, then fractionated using a HpH C18 column into 12 concatenated fractions as previously described (4).

**Myosin Heavy Chain Purification**

Myofibrillar protein purification was performed based on the method of (5). Tibialis anterior (TA) muscles from 10 wk old male C57Bl/6 mice were homogenised for 2 x 20 sec with an Omni Homogeniser (TH-02, Omni International, Kennesaw, GA, USA) in 2ml of buffer containing: 300 mM KCl, 10 mM Na4P2O7, 1.0 mM MgCl2, 150 mM Imidazole, 1.0 mM DTT, Protease inhibitor cocktail (P8340, Sigma, St Louis, MO, USA), pH 6.8. Samples were rotated for 90 min at 4oC and then centrifuged at 178,000 x g for 90 min at 4oC. The resulting supernatant was added to 32 ml of ice-cold H2O containing 1.0 mM DTT and protease inhibitor and incubated on ice for 60 min. After centrifugation at 20,000 x g at 4oC, the supernatant was carefully removed and the remaining pellet washed with cold H2O and then resuspended in 200 ul of storage buffer containing: 300 mM KCl, 1.0 mM EGTA, 4.0 mM MgCl2, 25mM Imidazole, 1.0 mM DTT and protease inhibitor cocktail, pH 7.3. Protein concentration was quantified using a DC protein assay kit (Bio-Rad, Hercules, CA). Proteins (6ug total) were then separated by SDS-PAE on a 4-20% gradient tris/glycine gel (Bio-Rad, Hercules, CA), and protein bands were visualized after Coomassie blue staining and destaining.

**LC-MS/MS data acquisition**

For analysis of mouse tissues, enriched and fractionated peptides were analyzed on a Dionex 3500 nanoHPLC coupled to an Orbitrap Exploris 480 mass spectrometer (Thermo Fisher Scientific) via electrospray ionisation in positive mode with 1.9 kV at 275 °C and RF set to 40%. Separation was achieved on a 50 cm × 75 µm column packed with C18AQ (Dr Maisch, Ammerbuch, Germany, 1.9 µm) (PepSep, Marslev, Denmark) over 54 mins at a flow rate of 300 nL/min. The peptides were eluted over a linear gradient of 3–25% Buffer B (Buffer A: 0.1% formic acid [FA]; Buffer B: 90% v/v acetonitrile, 0.1% v/v FA) and the column was maintained at 50 °C. The instrument was operated in data-dependent acquisition (DDA) mode with an MS1 spectrum acquired over the mass range 350–1,550 *m/z* (120,000 resolution, 300% automatic gain control (AGC) and 25 ms maximum injection time) followed by MS/MS analysis via HCD fragmentation mode and detection in the orbitrap (15,000 resolution, 1.2 *m/z* isolation, 200% AGC, 55 ms maximum injection time; 30% normalized collision energy). Fractionated TMT labelled enriched peptides from human skeletal muscle biopsies were analyzed on the identical chromatography conditions as described above except acquisition performed on an Orbitrap Ascend mass spectrometer (Thermo Fisher Scientific) via electrospray ionisation in positive mode with 2.0 kV at 275 °C and RF set to 30%. The instrument was operated in DDA mode with an MS1 spectrum acquired over the mass range 350–1,550 *m/z* (120,000 resolution, 250% AGC and 25 ms maximum injection time) followed by MS/MS analysis via HCD fragmentation mode and detection in the orbitrap (30,000 resolution, 1.2 *m/z* isolation, 200% AGC, 59 ms maximum injection time; 35% normalized collision energy, Enhanced Resolution Mode for TMTpro reagents).

**LC-MS/MS data processing**

Analysis of enriched peptides from various mouse tissues with SequestHT and Percolator (6) was performed in Proteome Discoverer (v2.5.0.4) against the UniProt mouse database (March 2022; UP000000589_10090 including isoforms with 63,592 entries). Data were filtered to 1% FDR at the peptide spectral match, peptide and protein level using QVALITY in the Protein FDR Validator node (7). All data were searched with oxidation of methionine (+15.994915), N-terminal protein acetylation (+42.010565), and VG of lysine (+156.08988) set as the variable modification. Carbamidomethylation of cysteine (+57.021464) was set as fixed modification. The MS1 mass tolerance was set to 10 ppm, while the mass tolerance for MS/MS fragments was set to 0.02 Da with maximum of 2 missed cleavage sites. Localisation of VG-sites was performed with PhosphoRS (8). Analysis of enriched peptides from various mouse tissues with MSFragger and PTMprophet was performed in FragPipe (v18.0) (9) against the identical database and same modifications described above. MSBooster (10) was enabled and data were filtered to 1% FDR at the peptide spectral match, peptide and protein level using ProteinProhet (11). Enriched and TMT-labelled peptides from mouse skeletal muscle treated with rAAV6 or human skeletal muscle biopsies were analysed with SequestHT and Percolator in Proteome Discoverer (v2.5.0.4) against the UniProt mouse database (March 2022; UP000000589_10090 including isoforms with 63,592 entries) or UniProt human database (March 2023; UP000005640_9606 including isoforms with 103,484 sequences), respectively. Data were searched with the parameters and modifications described above but also included TMTpro-VG of lysine (+460.297026) as a variable modification, and TMTpro of peptide N-terminus (+304.207146) as a fixed modification. Quantification was performed with the reporter ion quantification node for TMT quantification in Proteome Discoverer. TMT precision was set to 20 ppm and corrected for isotopic impurities. Only spectra with < 50% co-isolation interference were used for quantification.

**Quantification, statistical and downstream analysis**

For comparison of the 3 antibody clones and pooled cocktail, experiments were performed with one biological replicate. For comparison of mouse tissues, experiments were performed with one biological replicate. For comparison of mouse skeletal muscle treated with rAAV6:shScramble or rAAV6:shUFC1, experiments were performed with eight biological replicates each. For comparison of skeletal muscle from plwALS and age-matched controls, experiments were performed with eight biological replicates each. Data were processed with Perseus (12) with Log_2_-transformation and first normalized by subtracting the median of all the quantified non-VG-modified peptides of each sample to account of subtle differences in the amount of peptide enriched and labelled. The data were further normalized by subtracting the abundance of the protein to account of possible differences in the abundance of the VG-peptide vs protein levels. For comparison of mouse skeletal muscle treated with rAAV6:shScramble or rAAV6:shUFC1, statistical analysis was performed using two-way paired student’s t-tests and p-values correcting for multiple hypothesis testing using Benjamini Hochberg to obtain q-values. For comparison of skeletal muscle from plwALS and age-matched controls, statistical analysis was performed using two-way unpaired student’s t-tests and p-values correcting for multiple hypothesis testing using Benjamini Hochberg to obtain q-values. Significance thresholds are indicated in Figure Legends with p-value<0.05 or q-value<0.05.

**EvoEF protein stability simulations**

EvoEF Protein Stability Simulations EvoEF (version 1) was used to calculate the stability change upon mutation, in terms of ΔΔG. To this end, we first used "EvoEF --command=RepairStructure" to repair clashes and torsional angles of the wild type structure. "EvoEF --command=BuildMutant" is then used to mutate the repaired wild type structures into the mutant by changing the side chain amino acid type followed by a local side chain repacking. "EvoEF --command=ComputeStability" is then applied to both the repair wild type and the mutant to calculate their respective stabilities (ΔGWT and ΔGmutant). The stability change upon mutation can then be derived by ΔΔG=ΔGmutant-ΔGWT. A ΔΔG below zero means that the mutation causes destabilization; otherwise, it induces stabilization (13, 14). The sequence of the MYH2 coiled-coil backbone was used for EvoEF stability calculations due to size limitations in the software when using the entire MYH2 protein sequence.

**Data availability**

The mass spectrometry proteomics data have been deposited to the ProteomeXchange Consortium via the PRIDE (15) partner repository with the dataset identifier PXD051412 (Username:; Password: 3se7c5iG).

**Supporting Information References**
